# Phylogenetic analysis of Michigan’s freshwater sponges (Porifera, Spongillidae) using extended COI mtDNA sequences

**DOI:** 10.1101/2020.04.26.062448

**Authors:** Stephen H. Kolomyjec, Roger A. Willford, THE FALL 2019 GENETICS CLASS

## Abstract

Kolomyjec *et al.* (2020). Phylogenetic analysis of Michigan’s freshwater sponges (Porifera, Spongillidae) using extended COI mtDNA sequences. – *Zoologica Scripta*, 00, 000-000. The phylogenetic relationships of eight species of freshwater sponges sampled throughout the State of Michigan in the North American Great Lakes region were examined as part of a Course-based Undergraduate Research Experience (CURE). An extended version of the standard cytochrome oxidase subunit I (COI) metazoan DNA was used for sequencing and Bayesian phylogenetic inference. The extended gene region (COI-ext) produces a 1,200 bp amplicon instead of the standard 640 bp fragment which compensates for the standard amplicon’s low informatics value in Phylum Porifera. The species examined clustered into strongly supported monophyletic species groups within the family *Spongillidae*. This study represents the first look at the phylogenetic relationships of freshwater sponges in the Great Lakes Region.

Per Kolomyjec, College of Science and the Environment, Lake Superior State University, 650 W Easterday Ave., Sault Sainte Marie, MI 49783, USA.

## Introduction

Freshwater sponges are the pinnacle of the weird and wonderful. As a basal animal that lack some of the important aspects of the eumetazoans such a true tissues or organs, sponges are of keen interest to evolutionary biologists studying the origin and evolution of the multicellular animal body form (e.g. Nielsen 2008; Borchiellini *et al.* 2001). As sessile filter feeders, sponges provide a wide range of ecological roles and services from filtering water to providing habitat and refuge to countless organisms, even contributing to primary production in their local ecosystems through symbiosis with a wide variety of autotrophic organisms (Reiswig et al 2010; Manconi & Pronzato 2016). There is even the potential for extensive bioprospecting for the novel secondary metabolites formed by sponges and their extensive microbiome (e.g. Laport & Santos 2009; Simmons *et al.* 2005; Balasubramanian *et al.* 2018). Despite all of this, and the volume of research focused on marine sponges, most people are not even aware that freshwater sponges exist, and they remain relatively understudied.

The field of freshwater sponge research is a relatively small global community continually working towards an improved understanding of distribution and taxonomy at the continental to global scale (e.g. Manconi & Pronzato 2008, 2002, 2015; Reiswig *et al.* 2010; Ricciardi & Reiswig 1993; Volkmer-Ribeiro & Travest 1987). Of particular regional importance to the authors is the doctoral thesis and resulting publications of M. C. Old (1930, 1931) which detail the last and most complete state-wide survey of freshwater sponge diversity in Michigan. This past study places the reported number of species found in Michigan waters at 12 species within 10 genera (Old 1931), with a total of 31 species within 14 genera currently reported for continental North America (Manconi & Pronzato 2015).

The vast majority of research has been carried out using morphological features to determine species identification and taxonomy of the freshwater sponges. To date few studies have used molecular characters to explore freshwater sponge systematics and taxonomy (Addis & Peterson 2005). This dearth of information hampers research in freshwater biodiversity and conservation. Currently, identification of freshwater sponge species to the genus or species level requires the careful examination of microscopic traits based on spicule and gemmule morphology (e.g. Old 1931; Reswig *et al.* 2010; Manconi & Pronzato 2002). The seasonality of gemmules in many species and the high degree of phenotypic plasticity observed in freshwater sponges leads to the ability to meaningfully identify and quantify sponges to beyond the effort/benefit threshold of most freshwater research endeavors. This commonly leads to sponges only being included and quantified at the phylum level (MI DEQ 2000). This obviously renders a non-representative view of any system’s actual biodiversity and conservation needs.

A molecular approach to species identification, such as DNA barcoding, in biological research has the potential to alleviate this barrier and opens the possibility of species detection using aquatic eDNA approaches (e.g. Pilliod *et al.* 2013). However, this requires adequate baseline data which as mentioned is lacking for freshwater sponges. For example, at the time of this project GenBank (Sayers *et al.* 2019) only contained COI (Cytochrome Oxidase subunit I) sequences, commonly used for DNA barcoding of animals, for five of Michigan’s 12 species of sponges, with only three of those species having more than a single DNA record.

These gaps in the available data became evident during the first large scale resurvey of Michigan’s freshwater sponges in over 80 years (Old 1931) and the efforts reported in this study were focused on using the COI gene as a molecular marker for exploring the phylogenetic relationships of Michigan’s freshwater sponges and expanding the range of publicly available DNA sequences available to aide future research.

## Materials and Methods

### Course-based Undergraduate Research Experience (CURE)

This research was conducted as a course-based undergraduate research experience (CURE), which is an approach to integrating authentic research into the undergraduate curriculum in a manner that allows participation of a large number of students simultaneously (e.g. (Auchincloss *et al*. 2014; Rowland *et al*. 2012). This CURE was integrated into Dr. Kolomyjec’s 200-level genetics class in the fall 2019 semester at Lake Superior State University as an extension of the course’s regular DNA barcoding lab. There were 51 students actively involved across three lab sections. Individual students were tasked with DNA isolation, PCR and sequencing prep, sequence trimming and quality control, DNA based specimen identification (a.k.a. DNA barcoding) when possible, and initial phylogeny construction.

### Sponge samples

Fifty-one 70% ethanol preserved sponge specimens were semi-randomly selected from samples collected during a recent statewide survey (Willford & Kolomyjec unpublished). The only criteria for selection was that the individual specimen was large enough for students to subsample without difficult or risk of destroying the voucher specimen. The samples selected represented 17 of the 44 8-digit HUCs (Hydrologic Unit Codes) sampled by Willford & Kolomyjec (2019; unpublished).

### Taxonomic identification

Species level identification of freshwater sponges requires careful examination of the microscopic morphology of the hard-skeletal elements (spicules) and over-wintering structures (gemmules). The protocol used was modified from that of Old (1931). Subsamples of the selected specimens were placed in test tubes, dried overnight in a drying oven to remove all traces of ethanol and prevent an explosive reaction with the nitric acid used during tissue digestion. Tissue digestion was performed with concentrated nitric acid (HNO_3_) on a 108° C heat block in a fume hood until soft tissues of the sponge were completely dissolved. Nitric acid was then diluted with an equal volume of double-distilled water (ddH_2_O). After complete sedimentation of the spicules (about 10-15 minutes) as much liquid as possible was removed from the tube with a disposable transfer pipette. Fresh ddH_2_O was added and gently triturated with a new pipette to wash the spicules. Once again, the liquid was removed after sedimentation. The wash process was repeated at least 3 times. After the final wash, the pellet of spicules was resuspended in fresh ddH_2_O. A drop of suspended spicules was placed on a standard microscope slide and dried on the same heat block used during tissue digestion. Finally, slides were cover-slipped using a UV cured mounting medium (eukitt).

Species were independently identified by each of the authors using Old (1931), Reswig *et al.* (2010), and Manconi & Pronzato (2002, 2016) as primary references. Any IDs in disagreement were compared, reexamined, and resolved together. Genera specialist were consulted when additional confirmation was required.

### Nucleic acid extraction

Using aseptic techniques, students removed a portion of tissue approximately 3 mm × 3 mm from their assigned specimen for DNA isolation. DNA was isolated using a modified guanidine thiocyanate/silica resin extraction procedure (Cold Spring Harbor 2018). After samples were blot dried with lab tissues, they were placed in 1.5 ml microcentrifuge tubes with 300 μl 6M guanidine thiocyanate using disposable micro-pestles. The resultant slurry was incubated at 65° C for 10 min then centrifuged at 13,000 g for one minute to pellet out cell debris. 150 μl of the resulting supernatant was transferred to a clean 1.5 ml tube along with 3 μl of silica resin (aqueous 325 mesh SiO_2_, 50% w/v). This new tube was vortexed and incubated for 10 minutes at 57° C to bind silica and DNA. This was then pelleted by centrifugation at 13,000 g for 30 seconds. The supernatant was discarded, and the pellet washed twice with a wash buffer consisting of 1:1 mix of TNE buffer (1 M Tris (pH 7.4), 5 M NaCl, 0.5 M EDTA) and 100% Ethanol. The washed pellet was eluted with 100 μl Molecular Grade H_2_O. The unbound silica was pelleted out and the resultant supernatant transferred to a new tube for storage and downstream use.

DNA quality and quantity were estimated via spectrophotometry on a Bio-Spec Nano (Shimadzu).

### Polymerase chain reaction and sequencing

As the standard 640 bp animal barcoding fragment (Folmer *et al.* 1994) of the Cytochrome Oxidase I gene (COI) gene has been demonstrated to be of low informatics value in the sponges (Erpenbeck *et al.* 2006), a 1,200 bp extended version of the gene was amplified. This extended COI fragment was amplified using the dgLCO1490 (5’-GGT CAA CAA ATC ATA AAG AYA TYG G-3’) and COI-R1 (5’-TGT TGR GGG AAA AAR GTT AAA TT-3’) degenerate primers (synthesized by Invitrogen) from Meyer *et al.* (2005) and Rot *et al.* (2016) respectively. PCR reactions were performed in a 55 μl volumes containing: 25 μl Hot Start PCR-to-Gel Master Mix (VWR brand); 2 μl of primer mix (25 pmol each of forward and reverse primer); 5 μl template DNA; 23 μl PCR grade H_2_O. The thermal cycling regime included an initial denaturation phase of 95° C for 5 min followed by 30 cycles of: 40 sec denaturation at 95° C; 40 sec annealing 55°C; followed by 60 sec elongation at 72° C; and a final elongation at 72° C for 5 min. PCR success was checked on a standard 2% agarose gel visualized with GelRed nucleic acid stain (Biotium). Sequencing of successful PCR product was performed by the Psomagen Inc. (formerly MacrogenUSA) sequencing core using their LCO1490 universal primer.

### Phylogenetic analyses

A set of reference sequences (Table 1) was obtained from GenBank (Sayers *et al.* 2019) for comparison to the experimental samples. The marine demosponge *Tethya aurantium* was selected to serve as the outgroup for analysis. All sequences were trimmed, checked by amino acid reading frames to avoid frameshift errors in alignment, and finally aligned using BioEdit Version 7.0.5.3 (Hall 1999).

**Table 1.** COI reference sequences from genbank.

| **Species** | **Ascension #** |
| --- | --- |
| *Ephydatia fluviatilis* | JF340437.1 |
| *Ephydatia muelleri* | DQ176778.1 |
| *Eunapius fragilis* | DQ176779.1 |
| *Spongilla lacustris* | MH985292.1 |
| *Tethya aurantium\** | EF584565.1 |
| *Trochospongilla pennsylvanica* | DQ087503.1 |

*\*Marine demosponge selected as outgroup*

The program beast (v1.10.4; Suchard *et al*. 2018) was used for the Bayesian inference of phylogenetic relationships. The default HKY substitution model was utilized as it produced the best Bayesian information criterion score (BIC) in jModelTest v2.1.10 (Darriba *et al.* 2012). Markov chain Monte Carlo (MCMC) analyses were run for 1 billion generations (burn-in 200 million generations). Beast’s TreeAnnotator tool was used to estimate the posterior probability of inferred phylogenetic clades via the generation of a maximum clade credibility (MCC) summary tree. Trees were visualized using Beast’s FigTree (v1.4.4) tool and Archaeopteryx (v0.9928; Han & Zmasek 2009).

## Results

### Samples and sequencing

Students successfully isolated DNA from 51 freshwater sponge tissue samples. Of those 51 DNA isolates, 45 samples were successfully amplified via PCR of the COI locus. PCR product from three additional samples of interest, generated by the class instructor (S Kolomyjec) were included and 48 samples in total were sent off for Sanger sequencing. Of those 48 samples, 42 yielded high-quality sequence for downstream analysis.

While awaiting sequencing, authors Willford and Kolomyjec carried out independent morphological identifications of all specimens included in this study. Three samples (K19, K34, and K39) were initially listed as unknown due to a lack of gemmules which prevented conclusive differentiation between *Eunapius fragilis* and *Pottsiella aspinosa*. Identity of these samples was later confirmed via localization within the generated COI phylogeny.

### Extended Cytochrome Oxidase Subunit I (COI) mtDNA

All COI DNA sequences generated and analyzed in this study have been submitted to GenBank (Supplemental Table S1). Eight out of the twelve species reported to occur in Michigan (OLD 1931, 1932) were represented in this study. This adds COI sequences for three previously unrepresented species to the GenBank database: *Duosclera mackayi; Heteromeyenia baileyi; and Pottsiella aspinosa*.

When examining the maximum clade credibility (MCC) summary tree (Figure 1 & Supplemental Figure S2) generated by Beast’s Tree Annotator, 100% of samples clustered into monophyletic clades representing their species. These species clades were, in turn, supported by moderate to high posterior probability values (0.87-1.00) (Figure 1). Sister clades that support close relationships between species that have traditionally placed in close proximity due to morphological similarities are also present.

**Figure 1.**
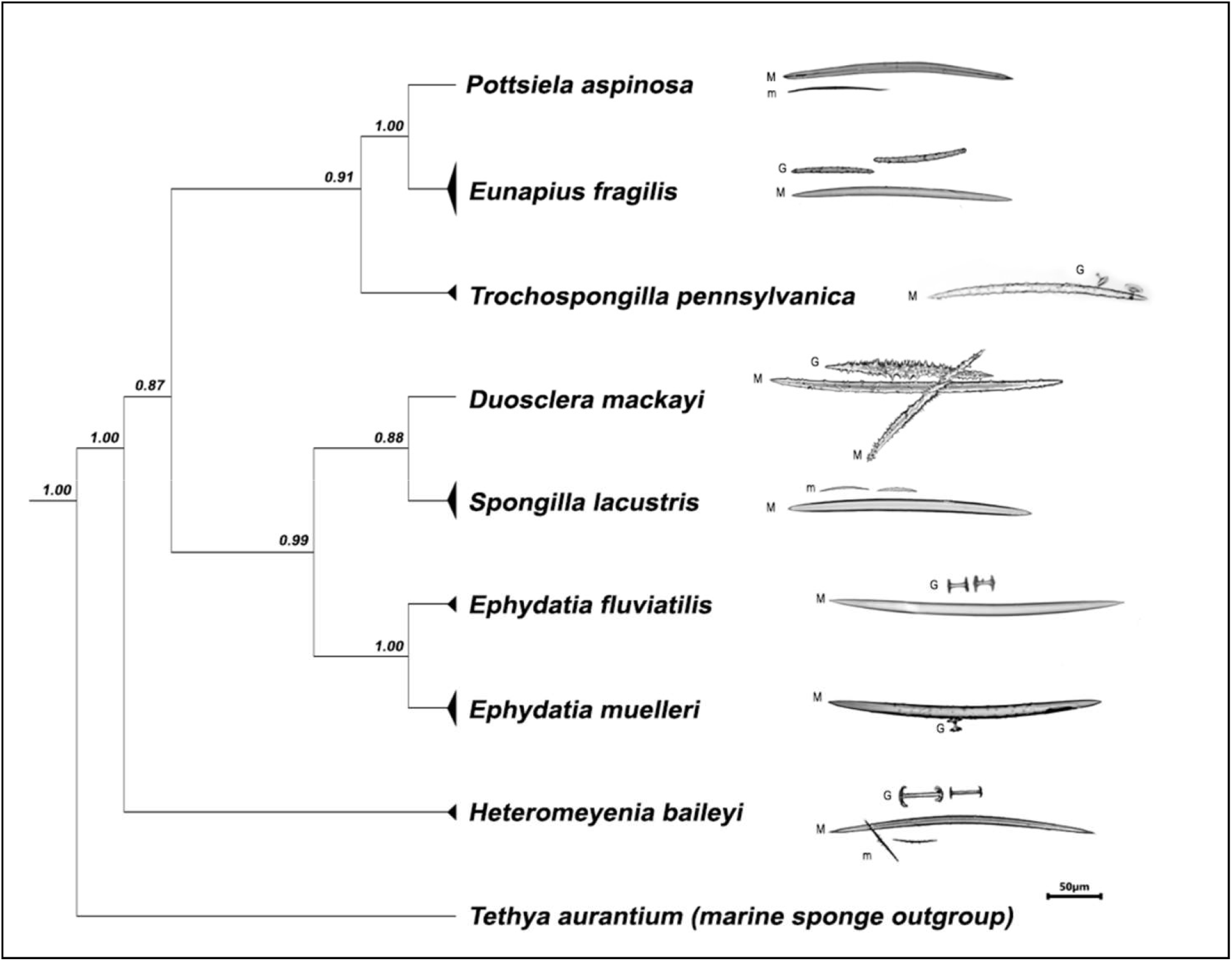
Relationship of Michigan’s freshwater sponge species. The summary cladogram depicts the relationships of species present in this study along with representative examples of spicules. Posterior probabilitie are indicated to the left of each node. Spicule coding: M = megasclere; m = megasclere; G = gemmulosclere.

## Discussion

### Phylogenetic Relationships of Michigan’s Freshwater Sponges

Analysis of an extended version of COI mtDNA (Erpenbeck *et al.* 2005) has provided a clearer phylogenetic interpretation of the relationships between freshwater sponge species than previous studies that utilized the standard COI segment (*e.g*. Addis & Peterson 2005; Meixner *et al*. 2007).This has allowed the authors to examine the interrelationships of Michigan’s freshwater sponges with a hereto unachieved clarity while still focusing on the region of DNA commonly used for DNA barcoding based species identification.

All species included in this study cleanly separated into clades matching their taxonomic identity (Figure 1). In the case of *Trochospongilla pennsyanica, Eunapius fragilis, Ephydatia fluviatilis, Ephydatia muelleri,* and *Spongilla lacustris* these clades included the appropriate species-specific reference sequences from GenBank. Intrageneric relationships are also maintained. For example, the two *Ephydatia* species present resolve as sister clades with a posterior probability of 1.00. In turn, the inclusive *Ephydatia* clade is sister to a larger clade containing *S. lacustris* and *Duosclera mackay*i. Interestingly enough *D. mackayi* was originally included within the *Spongilla* genera until Reiswig and Ricciardi (1993) placed it in the monotypic genera *Duosclera* on the basis of extensively unique morphology.

We were able to comfortably assign the three samples that were inconclusively identified via standard light microscopy (K19, K34, and K39) to the *Eunapius fragilis* clade based on their unambiguous phylogenetic placement (S2). Considering the difficulty inherent in identifying freshwater sponges to the species level, this is no minor accomplishment. Setting aside phenotypic variation that often results from the same species of sponge growing under slightly different ecological variation, not all individual sponges will produce all the structures required for definitive identification at all stages of their life cycle. This is a well-established issue in freshwater sponge research (Reiswig et al 2010; Manconi & Pronzato 2016) that can lead to the necessity of repeated sampling across both time and space to resolve the identity of a single species of freshwater sponge within a body of water. Considering the time and monetary expenses of fieldwork in general, the development and validation of supplementary tools such a DNA based identification and verification is critical to the consistent and reliable reporting of species level biodiversity. This study represents a major step forward in refining these tools for freshwater sponge research. While the primary focus of this study was regionally specific to Michigan and the Great Lakes region, many of the species present have a wider, or even global, distribution.

While this study provides vital reference sequences and helps to expand the knowledge of the relationships of the freshwater sponges in Michigan and to a lesser extent globally, there is much to be done within the field as a whole. Our research group will continue to work on expanding the sequence-based representation of regional species but to truly understand the relationships across all freshwater sponges a great deal more work needs to be done at the continental and global scale. This work should utilize additional genes or ideally whole genome comparison for hierarchical reconstruction of hydrologically influenced population structure and true global phylogeny.

## Acknowledgements

We would like to recognize the students of the Fall 2019 Genetics Class at Lake Superior State University for their hard work diligence in the DNA isolation and generation of the raw sequence data used in this research: Emily Aisthorpe; Allie Anderson; Allison Blank; Haven Borghi; Kyle Burton; Veronica Clark; Nicole Clinard; Robert Cooper; Devon DePauw; Cameron Evans; Michael Gills; Tristen Gleason; David Gray; Michael Gray; Julianne Grenn; Wayra Hernandez-Mendez; Brady Hess; Kyra Hill; Olivia Hohman; Keegan Hoose; Brett Immel; Samuel Johnston; Callie Kammers; Ella Knopp; Alston Krikorian; Matthew Kurin; Rebecca Lathrop; Robert Lawrence; Ashley Lolmaugh; Isaac Loutzenhiser; Madison Marsh; Marissa McIntyre; Hayley Mellon; John Miles; Emily Miller; Brendan Montie; Lyndsey Murdock; Andrew Niemiec; Stephanie Owens; Sophia Passino; Brianna Regan; Brad Renner; Carly Sanofsky; Trenton Schipper; Courtney Simpson; Lindsay Smeltzer; Angelina Stout; Katie Thompson; Hunter VeltKamp; Rebecca Weipert; Gary Xiong.

We would also like to thank Lake Superior State University for the facilities, instrumentation, and logistics that made this project possible. We would like to specifically acknowledge Dr. Steven Johnson, dean of the College of Science and the Environment for continually supporting student-faculty research collaboration.

Finally, we would like to support experiment.com and all of the generous donors that helped fund the field collection of our freshwater sponge samples.

---

**Table S1.**
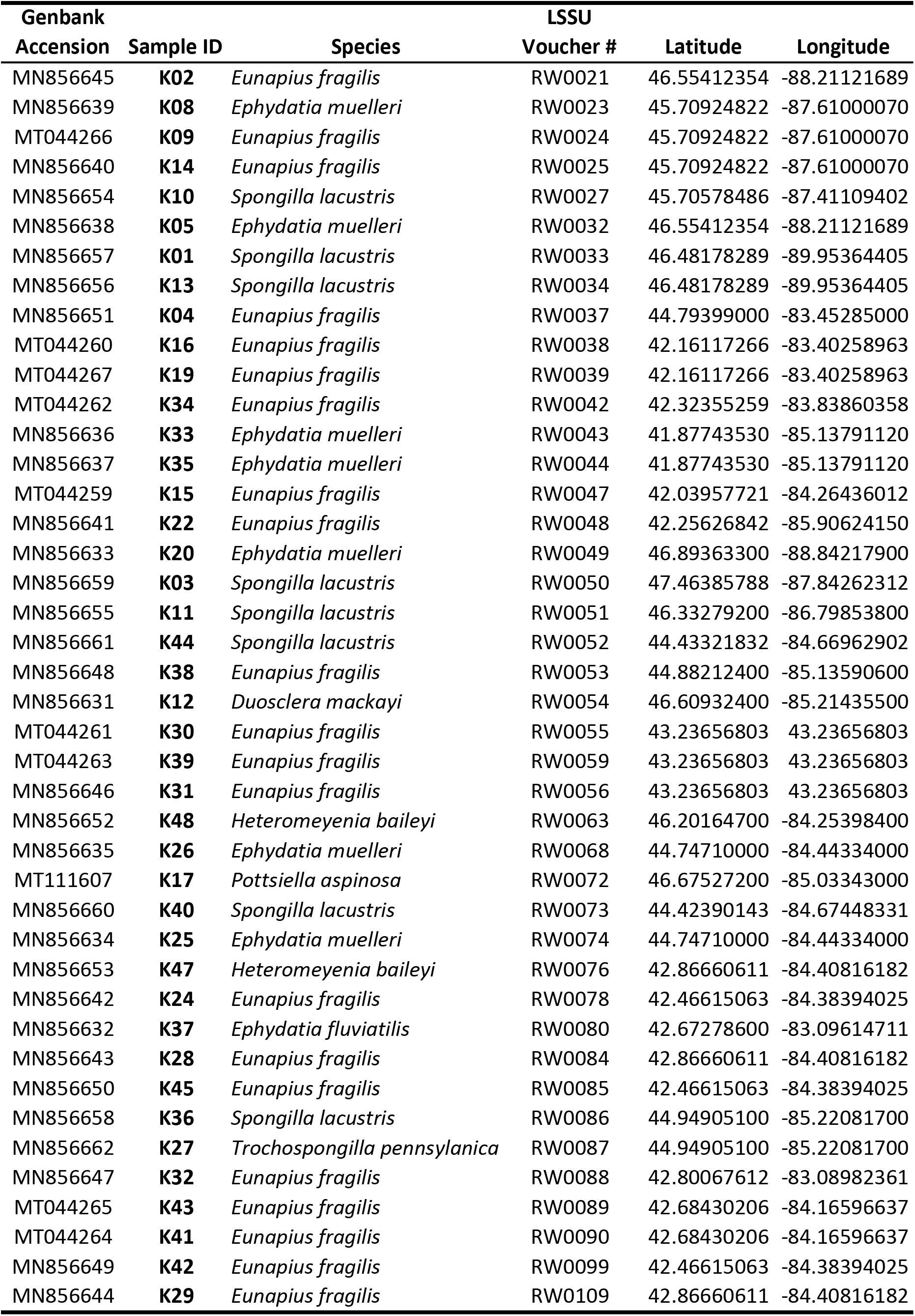
Genbank accension numbers and sample location for of sequences generated during this study.

**Supplemental Figure S2.**
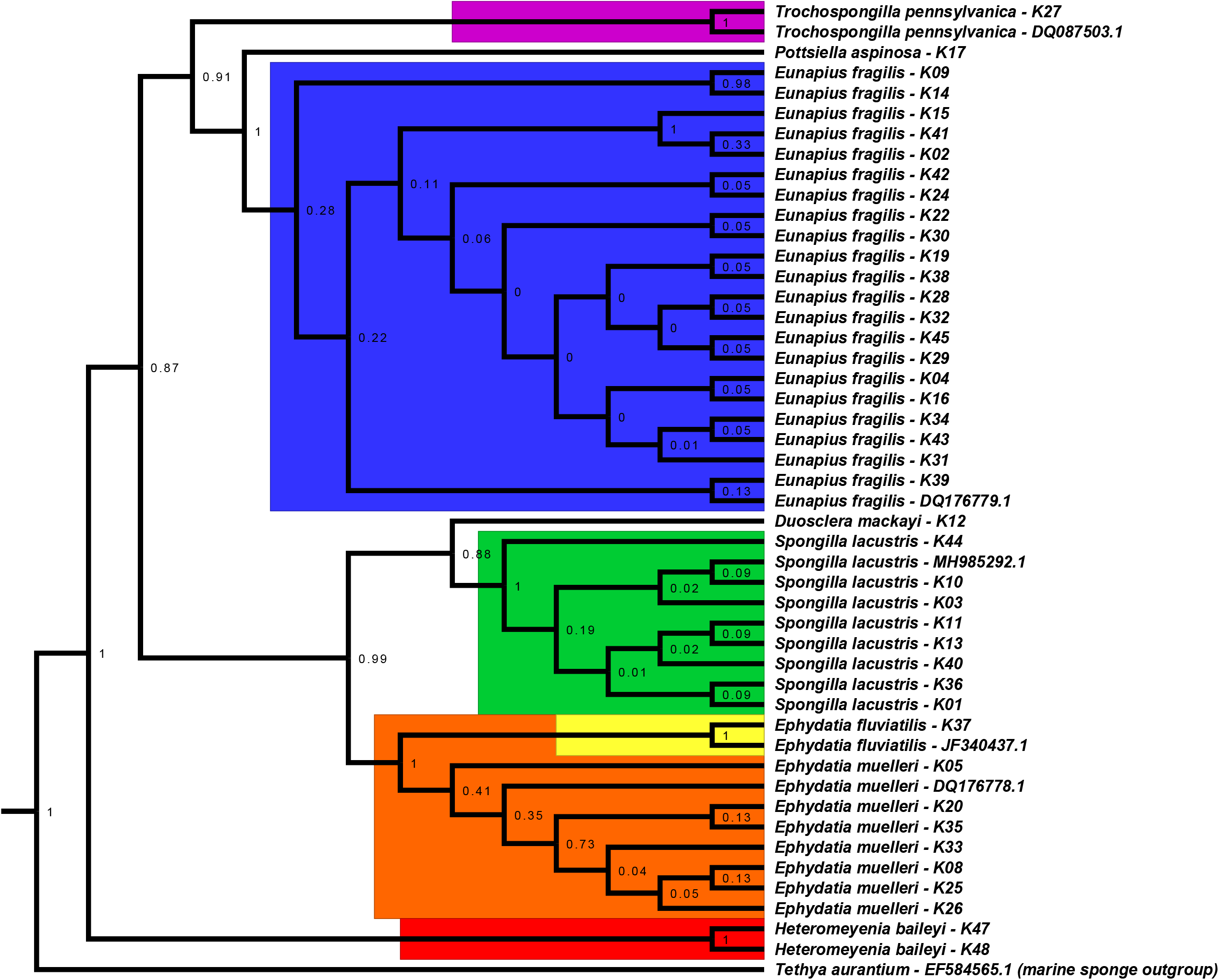
Full cladogram of all samples and reference sequences analysed during this study. Species clades are color coded for emphasis.

